# Designing AAV Capsid Protein with viability-guided Diffusion Model

**DOI:** 10.64898/2026.01.12.698995

**Authors:** Shuyuan Xiao, Xi Zeng, Shaoqing Jiao, Dazhi Lu, Dongna Xie, Jiaming Liu, Jiajie Peng

## Abstract

Adeno-associated virus (AAV) capsids have shown great promise as delivery vectors for gene therapy. However, the natural properties of AAV capsids impose significant limitations on their efficacy. While capsid library screening and directed evolution have facilitated the development of improved vectors, current methods for generating AAV libraries frequently yield nonviable variants that fail to assemble or package DNA. Here, we propose a viability-guided diffusion model AAVDiffusion for de novo viable AAV capsid design. AAVDiffusion iteratively denoises Gaussian vectors into vectors of AAV capsid protein sequences, yielding intermediate latent variables. By leveraging the continuous nature of these latent variables, AAVDiffusion integrates an additional viability classifier that applies gradient updates to enhance the generation of viable AAV sequences. Through extensive computational tests, AAVDiffusion exhibits superior performance in generating viable AAV sequences. Furthermore, 196 AAV candidates were identified through a selection workflow, with the potential to cross the blood-brain barrier, thereby offering safer and more effective gene therapies for brain diseases. AAVDiffusion offers a powerful and efficient computational method for designing viable AAV capsids, advancing the development of AAV vectors for gene therapy.

## 1 Introduction

Gene therapy involves delivering genetic material to patients to correct, augment, or inhibit a biological function for the treatment of diseases [1]. The success of such treatments largely depends on the efficiency and specificity of the delivery vehicle. Adeno-associated virus (AAV) vector is one of the most widely used vectors for gene therapy [2, 3]. AAV-based gene delivery vehicles are achieving growing success in human clinical trials, promising to treat a wide range of genetic and non-genetic diseases [4–11]. However, the current generation of AAVs remains insufficient to safely and effectively address the majority of human diseases. To expand their therapeutic potential, there is a critical need to develop new AAV variants with enhanced transduction efficiency, improved organ specificity, and reduced immunogenicity [12]. The key properties of AAV vectors are primarily determined by their capsid proteins [13], making the rational design of novel capsid proteins essential for advancing AAV vector engineering.

AAV capsid library selection and directed evolution have facilitated the development of significantly improved vectors, some of which have progressed into clinical applications [14, 15]. A fundamental criterion for the success of capsid library generation is the viability of capsid variants, ensuring that engineered AAV capsids can efficiently assemble and package the viral genome [16–18]. Unfortunately, many widely used methods for generating AAV capsid libraries—such as error-prone mutagenesis [19], random shuffling between AAV serotypes [20], and peptide display [21]—produce a high proportion of nonviable variants. This highlights an urgent need for more efficient strategies to generate viable AAV capsid sequence variants, thereby accelerating the development of next-generation gene therapies.

In recent years, machine learning (ML) has been applied to viable AAV capsid design by developing predictive models. These models predict AAV viability based on sequence information and can be extended to generative methods by incorporating random AAV sequence generation. For example, Bryant et al. utilized logistic regression (LR), convolutional neural networks (CNNs), and recurrent neural networks (RNNs) to predict the viability of AAV capsid sequence variants [22]. Moreover, Eid et al. developed a long short-term memory (LSTM) regression model to predict the viability of AAV-7mer variants, leveraging LSTM’s ability to capture local and distant relationships across different parts of the input sequences [23]. Marques et al. trained artificial neural networks (ANNs) and support vector machines (SVMs) to predict the viability of AAV capsid sequence variants [24]. Despite these advancements, the inherent scarcity of viable variants within the vast protein sequence space leads to inefficient sampling in these models, ultimately limiting their predictive performance and design efficiency. In contrast to predictive models, generative models can learn the underlying data distribution and directly generate novel protein sequences, providing efficient computational approaches for de novo protein design. Commonly used generative frameworks include Autoregressive Models (AMs) [25–27], Variational Autoencoders (VAEs) [28, 29], and Generative Adversarial Networks (GANs) [30], all of which have demonstrated success in generating functional proteins [31–41]. However, despite the widespread application of generative models in protein design, their potential for viable AAV capsid engineering remains largely underexplored.

To improve the efficiency of generating viable AAV capsid proteins, we introduce AAVDiffusion, a viability-guided diffusion model specifically designed for the de novo generation of viable AAV capsid sequences. The model encodes AAV protein sequences into a latent space and progressively introduces Gaussian noise during the forward diffusion process, gradually perturbing their structure [42]. In the denoising phase, the model iteratively predicts and removes noise through a learned reverse process, ultimately reconstructing viable protein sequences from their latent representations. Unlike conventional diffusion models that employ U-Net architectures for image processing [43], AAVDiffusion leverages a Transformer-based architecture for denoising, utilizing the self-attention mechanism to capture long-range dependencies and structural information in protein sequences. To enhance its ability to learn the underlying “grammar” of functional peptides and expand biological diversity during training, we pre-train the model on peptide sequences from the UniProt database [35, 37, 44]. In addition, for precise control over the viability of generated sequences, AAVDiffusion employs a lightweight, modular plug-and-play strategy [45], in which a classifier is trained on diffusion latent representations. During generation, gradient updates are applied to these latent representations, guiding the model toward producing viable AAV capsid sequences.

Following the existing work [22], we select the 561-588 segment of the AAV serotypes 2 (AAV2) VP1 protein for generation, a region of immunological significance that overlaps with both heparin– and antibody-binding sites [46]. With AAVDiffusion, we can generate a series of viable AAV capsid variants for screening enhanced properties. Computational evaluations of AAV capsid candidates indicate that AAVDiffusion outperforms other state-of-the-art methods [22, 34, 37, 39]. In addition, we identified 196 AAV capsid variants with higher binding affinities to the human transferrin receptor (hTfR1) than BI-hTfR1 [47], a recently designed AAV variant engineered to efficiently cross the blood-brain barrier (BBB) [48–52] via hTfR1 binding. These capsid variants show potential to cross the BBB, target neural cells, and provide safer, more effective gene therapies for brain diseases. Given the performance demonstrated in the evaluations, AAVDiffusion provides a powerful computational method for generating viable AAV capsids, thereby advancing the development of AAV vectors in gene therapy.

## 2 Results

### 2.1 AAVDiffusion – a viability-guided diffusion model for AAV capsid design

To de novo design AAV capsid proteins meeting the desired viability, we propose AAVDiffusion (Fig. 1), which comprises two major modules: AAV sequence diffusion module (Fig. 1a) and viability classifier module (Fig. 1b).

**Fig. 1.**
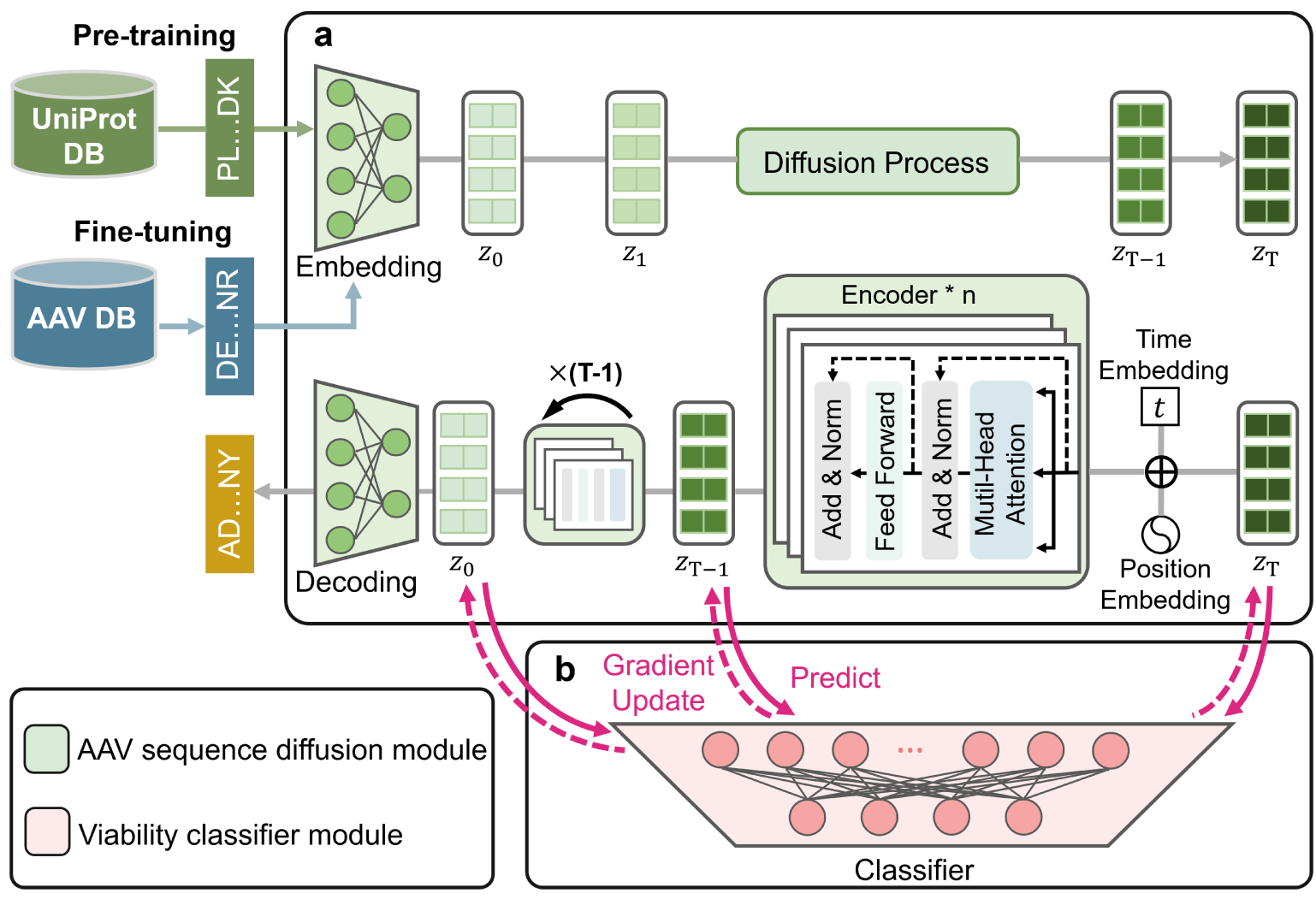
Overview of AAVDiffusion. **a**, AAV sequence diffusion module. The module consists of an embedding layer, a latent diffusion model and a decoding layer. The embedding layer is utilized to convert AAV sequences into low-dimensional latent representations *z*_0_. The latent diffusion model is designed to generate a new latent representation *z*_0_. The model consists of a diffusion process and a denoising process. The diffusion process adds Gaussian noise incrementally over *T* steps, trans-forming *z*_0_ into a noisy state *z_T_*. A Transformer-based denoising model is then trained to predict the noise added at each step. In the denoising process, starting from *z_T_*, the denoising model progressively denoises the latent vector to reconstruct *z*_0_. The decoding layer is utilized to reconstruct AAV sequence from the latent representation *z*_0_. **b,** Viability classifier module. The classifier ensures precise control over the viability of generated AAV sequences. It uses the latent representations of AAV sequences to predict sequence viability and guides the sequence generation toward high-viability designs by applying gradient updates to the latent variables at each denoising step.

The AAV sequence diffusion module aims to generate diverse AAV sequences by learning to maximize the marginal likelihood of the AAV sequence data. It was trained on the AAV dataset constructed by merging three datasets from Branyt et al. [22]. In this module, AAV sequences are first fed into the embedding layer to obtain low-dimensional latent representations *z*_0_, capturing key features of these proteins. During the diffusion process, starting with the latent vector *z*_0_, Gaussian noise is progressively added over *T* discrete steps via a Markov process *q*(*z_t_*|*z_t−_*_1_), culminating in a highly corrupted state *z_T_*. Then, a Transformer-based denoising network *f_θ_* is trained to predict the noise added at each step [27], effectively parameterizing the reverse Markov process *p_θ_*(*z_t−_*_1_|*z_t_*). In the denoising process, beginning with Gaussian noise *z_T_*, the denoising network progressively denoises the latent vector to reconstruct *z*_0_. Finally, the decoding layer maps the reconstructed *z*_0_ back to the protein sequence space, generating a novel AAV sequence. To better capture the “grammar” of functional peptides and enhance biological diversity, the AAV sequence diffusion module was pre-trained on peptide sequences from the UniProt database [44].

The viability classifier module is designed to ensure precise control over the viability of generated AAV sequences. After training the diffusion module, we trained a binary viability classifier on the latent representations of AAV sequences. Specifically, AAV sequences with binary viability labels are fed into the embedding layer to obtain their latent representations *z*_0_, which undergo the diffusion process to generate a series of latent vectors *z*_1:_*_T_*. The complete set of latent vectors *z*_0:_*_T_*, paired with their viability labels, was used to train the classifier to predict the viability of AAV sequences. In the controllable generation, the classifier performs gradient updates on the latent variable at each denoising step, guiding it toward high viability. This iterative process balances two objectives: maintaining high data likelihood and achieving high viability. Finally, the denoised latent representation *z*_0_ by gradient-guided is fed into the decoding layer to produce a viable AAV sequence. Further details about the AAVDiffusion can be found in the Methods section.

### 2.2 AAVDiffusion outperforms other models in generating viable AAV capsid variants

To evaluate the performance of AAVDiffusion in generating viable AAV sequence variants, we compared it with four state-of-the-art methods: ClaSS [37], MSA-VAE [34], Bryant et al.’s Method (referred to as DeepAAV) [22], and ProteinGAN [39]. These models generated 19,680 unique viable sequences, which were then evaluated using six viability classifiers developed by Bryant et al. [22], including CNN(C1+R2), CNN(C1+R10), CNN(R10+A39), RNN(C1+R2), RNN(C1+R10), and RNN(R10+A39). These classifiers are CNN and RNN models trained on three distinct datasets: C1+R2, C1+R10, and R10+A39. Each classifier predicts a viability probability for a given sequence, where a higher value suggests a greater likelihood of the sequence being viable for DNA payload packaging. AAVDiffusion achieved the highest median viability probability across all six classifiers, outperforming MSA-VAE, DeepAAV, ProteinGAN, and ClaSS. Particularly, under the evaluation of CNN(C1+R2), the median viability probability of AAVDiffusion surpasses that of the second-best model by 0.19 (Fig. 2a). These results demonstrate the superior ability of AAVDiffusion to generate viable AAV sequences.

**Fig. 2.**
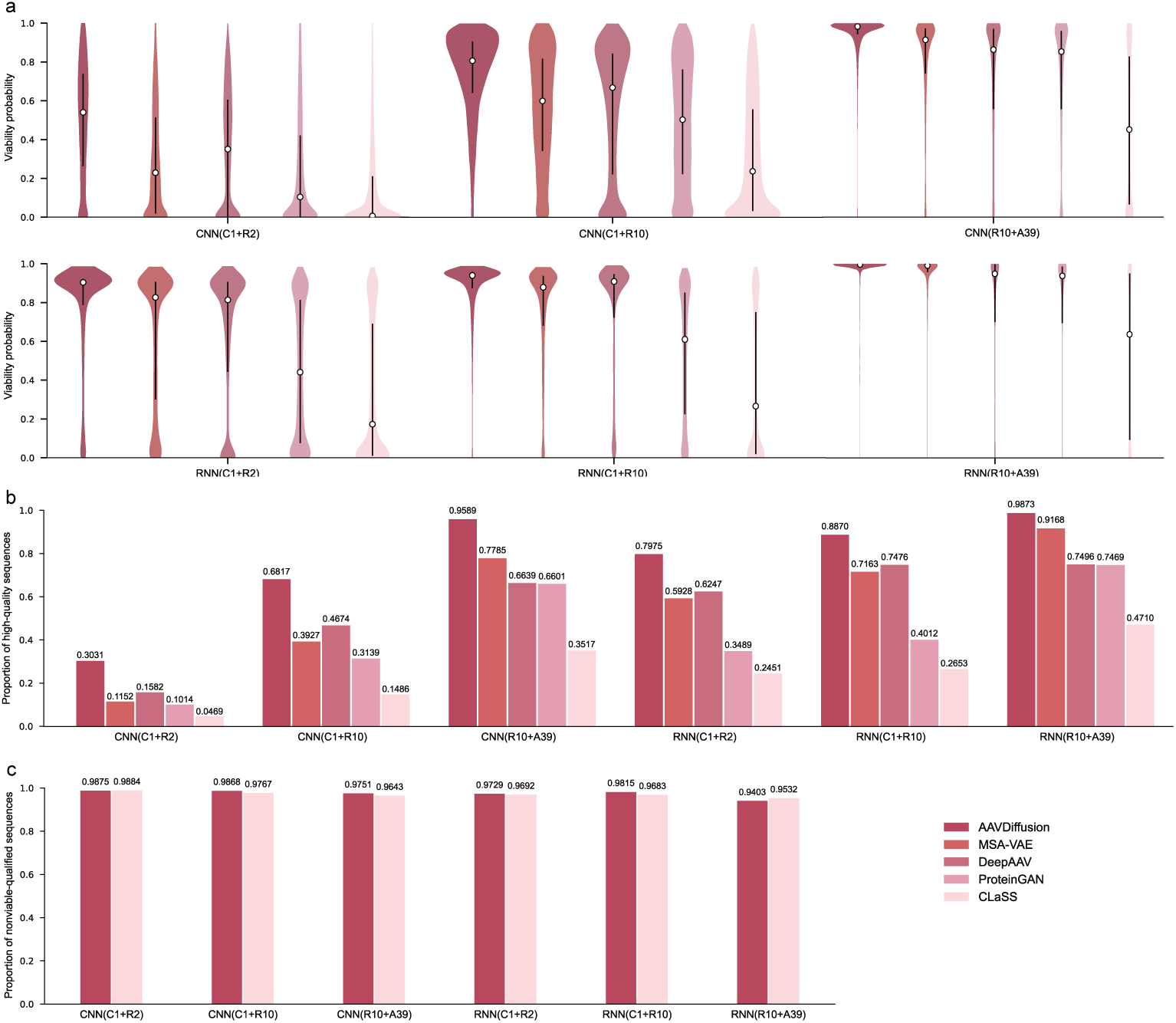
Controllable generation performance of AAVDiffusion, in comparison to MSA-VAE, DeepAAV, ProteinGAN, and ClaSS. **a**, Distributions of viability probability for viable sequences generated by AAVDiffusion and the other methods. The white dots mark the median of each distribution and the black vertical lines denote the interquartile range of each distribution. **b,** Proportions of high-quality sequences for AAVDiffusion and the other methods. **c,** Proportions of nonviable-qualified sequences for AAVDiffusion and ClaSS.

To further assess the ability of AAVDiffusion to generate high-viability sequences, we defined a classification threshold, where sequences with a viability probability *>* 0.7 were classified as high-quality, while others were classified as low-quality. Through the classification threshold, we found that AAVDiffusion achieved the highest proportion of high-quality sequences across all classifiers, with proportions of 30.31%, 68.17%, 95.89%, 79.75%, 88.70%, and 98.73% (Fig. 2b). These results demonstrat that AAVDiffusion outperforms the other methods in generating high-viability AAV sequences.

To evaluate the robustness of AAVDiffusion, we compared its generated 19,680 nonviable sequences with those generated by ClaSS, which can generate nonviable sequences and is resistant to false-negative predictions [37]. Sequences with a viability probability 0.5 were classified as nonviable-qualified, while others were considered viable-qualified. The results show that AAVDiffusion achieves a comparable propor-tion ( 95%) of nonviable-qualified sequences across all classifiers, demonstrating its capability to generate nonviable sequences. Overall, all these results highlight the strength of AAVDiffusion in generating viable AAV sequences.

### 2.3 The effects of pre-training and viability classifier

To assess the contributions of pre-training and viability classifier to the performance of AAVDiffusion, we conducted comprehensive ablation studies using four model configurations: (1) the AAVDiffusion model; (2) AAVDiffusion without pretraining; (3) AAVDiffusion without classifier; and (4) AAVDiffusion without both pretraining and classifier. All models generated 19,680 viable and 19,680 nonviable sequences, with sequence viability evaluated using the six classifiers mentioned above and the same classification thresholds.

For viable sequence generation (Figs. 3a, b), compared to AAVDiffusion without pre-training and classifier, adding pre-training increased median viability probability by 0.136 and raised the proportion of high-quality sequences by 7.81% on average. This suggests that pre-training helps AAVDiffusion better capture structural and functional constraints inherent to viable sequences by learning biologically meaningful patterns from the UniProt dataset, thereby improving its ability to generate high-quality sequences. Similarly, introducing the classifier increased the median viability probability by 0.232 and the proportion of high-quality sequences by 29.74% on average, demonstrating the gradient-based optimization of the classifier effectively improved sequence viability. When pre-training and the classifier were applied together, performance improved further, confirming that the combination significantly enhanced the generation of viable sequences.

**Fig. 3.**
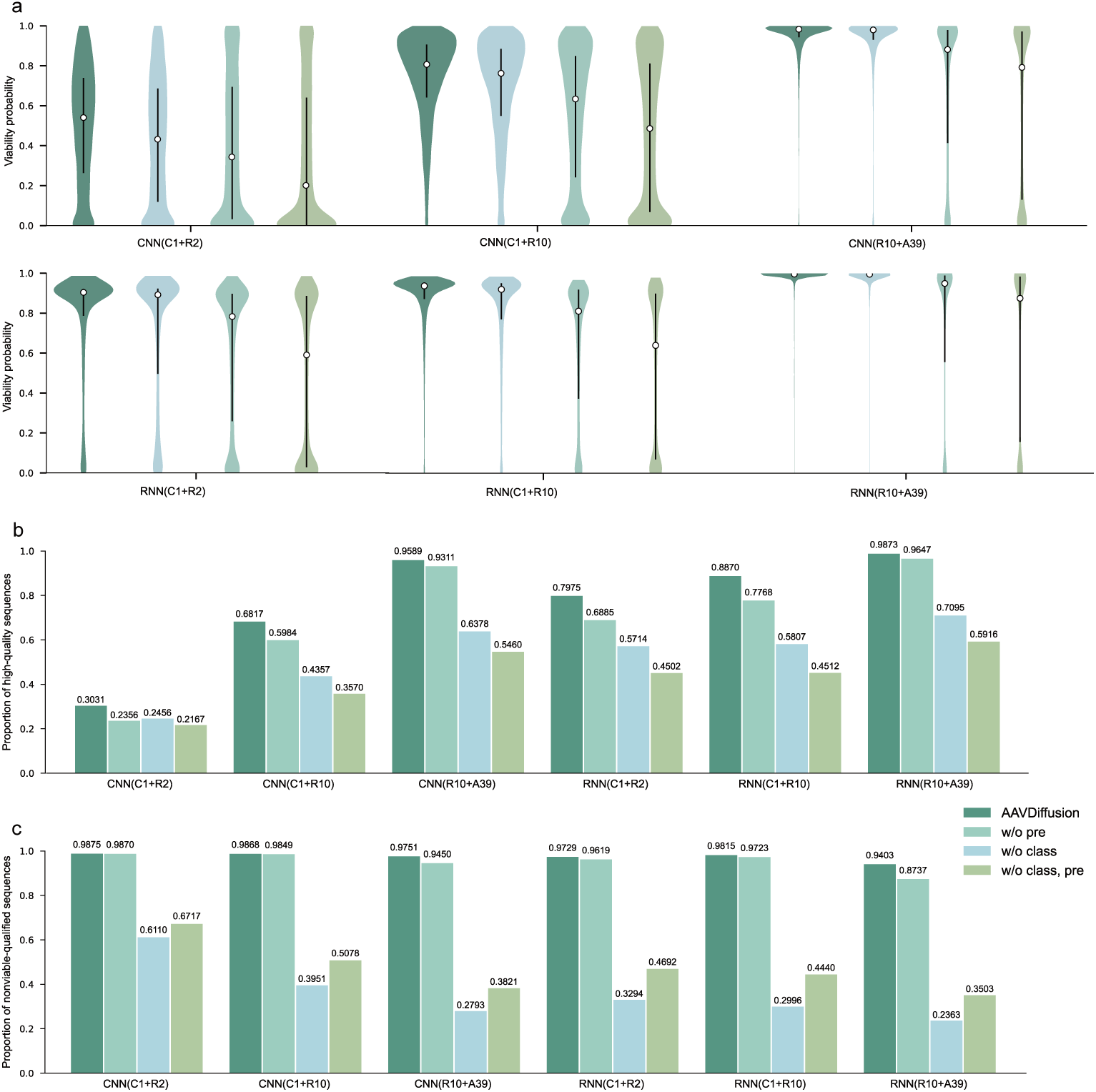
Ablation studies for AAVDiffusion. **a**, Distributions of viability probability for viable sequences generated by the AAVDiffusion model (AAVDiffusion), AAVDiffusion without pre-training (w/o pre), AAVDiffusion without classifier (w/o class) and AAVDiffusion without both pre-training and classifier (w/o class, pre). **b,** Proportions of high-quality sequences for the four model configura-tions. **c,** Proportions of nonviable-qualified sequences for the four model configurations.

For nonviable sequence generation (Fig. 3c), similar results were observed. Compared to the model without pre-training and classifier, adding pre-training increased the proportion of nonviable-qualified sequences by 48.33%. Incorporating both pre-training and the classifier further increased this proportion by 50.30%, emphasizing the combined effect on the robustness of AAVDiffison. When both components were removed, the proportion of nonviable-qualified sequences was higher than the configuration without the classifier alone. This suggests that in the uncontrolled generation, pre-training enhances sequence viability, thereby reducing the number of sequences classified as nonviable-qualified. Overall, these results highlight the critical roles of pre-training and viability classifier in AAVDiffusion and demonstrate that their combination significantly improves the model’s ability to generate viable sequences.

### 2.4 AAVDiffusion learns the intrinsic relationships of viable AAV sequences

To assess the capability of AAVDiffusion to capture the key features influencing AAV production, we analyzed the correlation between the viability probabilities predicted by the AAVDiffusion-Classifier and the experimental production scores. These scores were obtained from high-throughput experiments and quantify the efficiency of each protein sequence in generating viral particles [22]. We first obtained the latent representations *z_t_* of all sequences from the AAV dataset and used the AAVDiffusion-Classifier to predict the viability probabilities for each representation. Then the Pearson correlation coefficient between these predictions and experimental production scores was computed to quantify their correlation. The result shows a strong positive correlation (Pearson correlation coefficient = 0.78*, P <* 1 10*^−^*^16^) between them, indicating that AAVDiffusion effectively captures the critical features underlying AAV production (Fig. 4a).

**Fig. 4.**
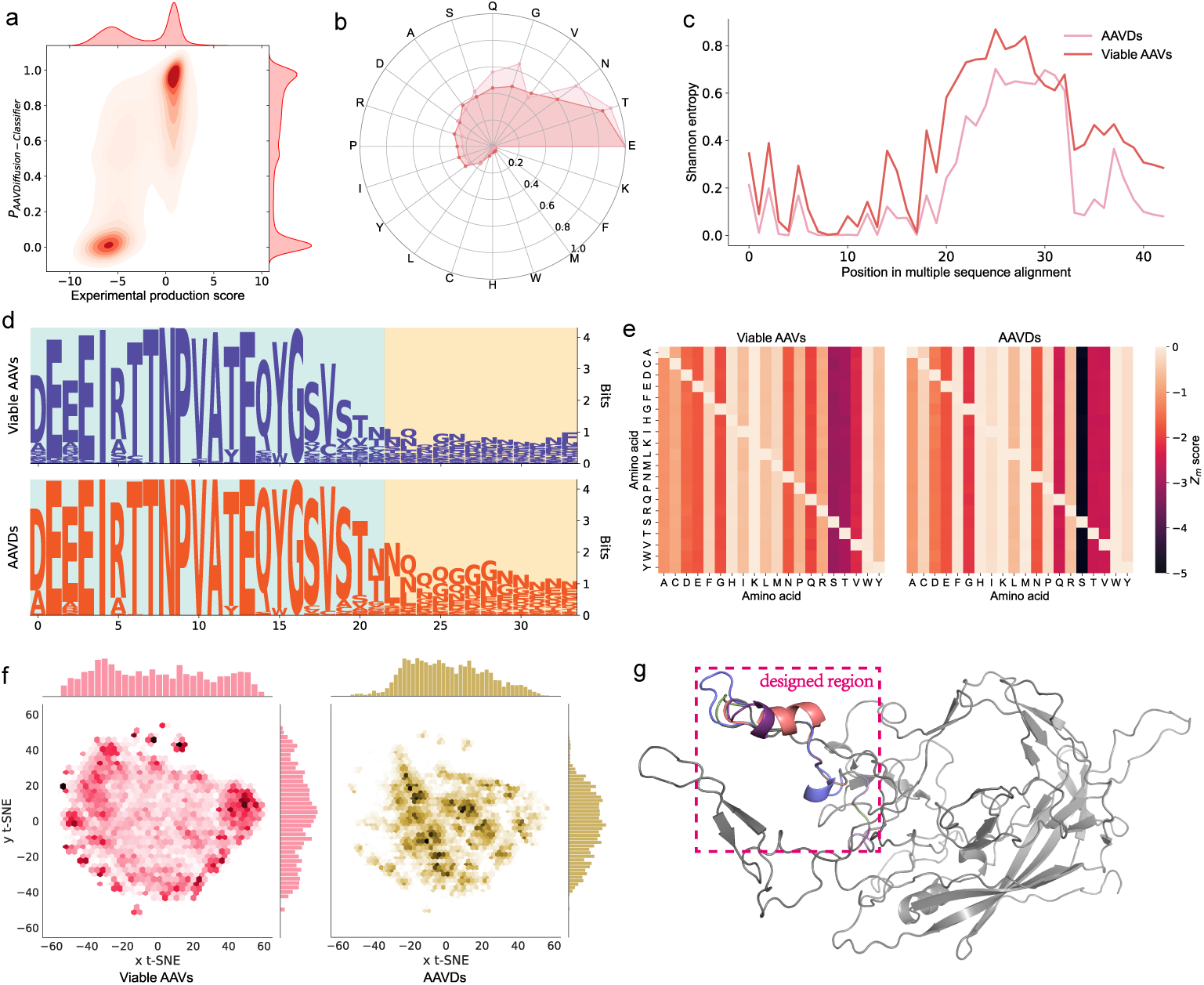
AAVDiffusion learns the intrinsic relationships between viable AAV sequences. **a**, Correlation between viability probabilities predicted by AAVDiffusion-Classifier and experimental production scores. **b**, Comparison of amino acid composition in Viable AAVs and AAVDs. **c**, Com-parison of Shannon entropies for Viable AAVs and AAVDs estimated from MSA. **d**, Sequence motif logos for Viable AAVs and AAVDs. **e**, *Zm* positional score matrices for Viable AAVs and AAVDs. **f**, t-SNE visualization of Viable AAVs and AAVDs. The color intensity of each dot represents data density, with darker colors indicating higher densities. **g**, Structural alignment of four AAVDs with the WT structure. The pink region is the designed region of the AAV2 VP1 sequence. Colored struc-tures denote the four generated sequences, while the gray represents the WT sequence.

To evaluate the ability of AAVDiffusion to capture the fundamental molecular features of viable AAV sequences, we compared the amino acid composition of 19,680 randomly selected viable sequences from the AAV dataset (labeled as Viable AAVs) with 19,680 AAV sequences generated by AAVDiffusion (labeled as AAVDs). The analysis reveals a high degree of consistency in the overall amino acid composition between Viable AAVs and AAVD, despite specific compositional variations. Specifically, AAVDs are relatively more enriched in Gly, Asn, and Gln, while showing a relative depletion of Ile, Pro, Arg, and Tyr (Fig. 4b). Furthermore, we analyzed the most abundant k-mers (with *k* = 3, 4, 5) in both datasets (Sup. Table 1), revealing a shared preference for k-mers rich in Thr, Asn, Asn, Pro, and Val. Compared to Viable AAVs, AAVDs exhibit lower frequencies of the highest-frequency k-mers, indicating greater sequence diversity. These results demonstrate that AAVDiffusion effectively captures the compositional and motiflevel features of Viable AAVs. Moreover, it introduces distinct compositional shifts and enhances sequence diversity, highlighting its potential for novel AAV sequence design.

**Table 1.**
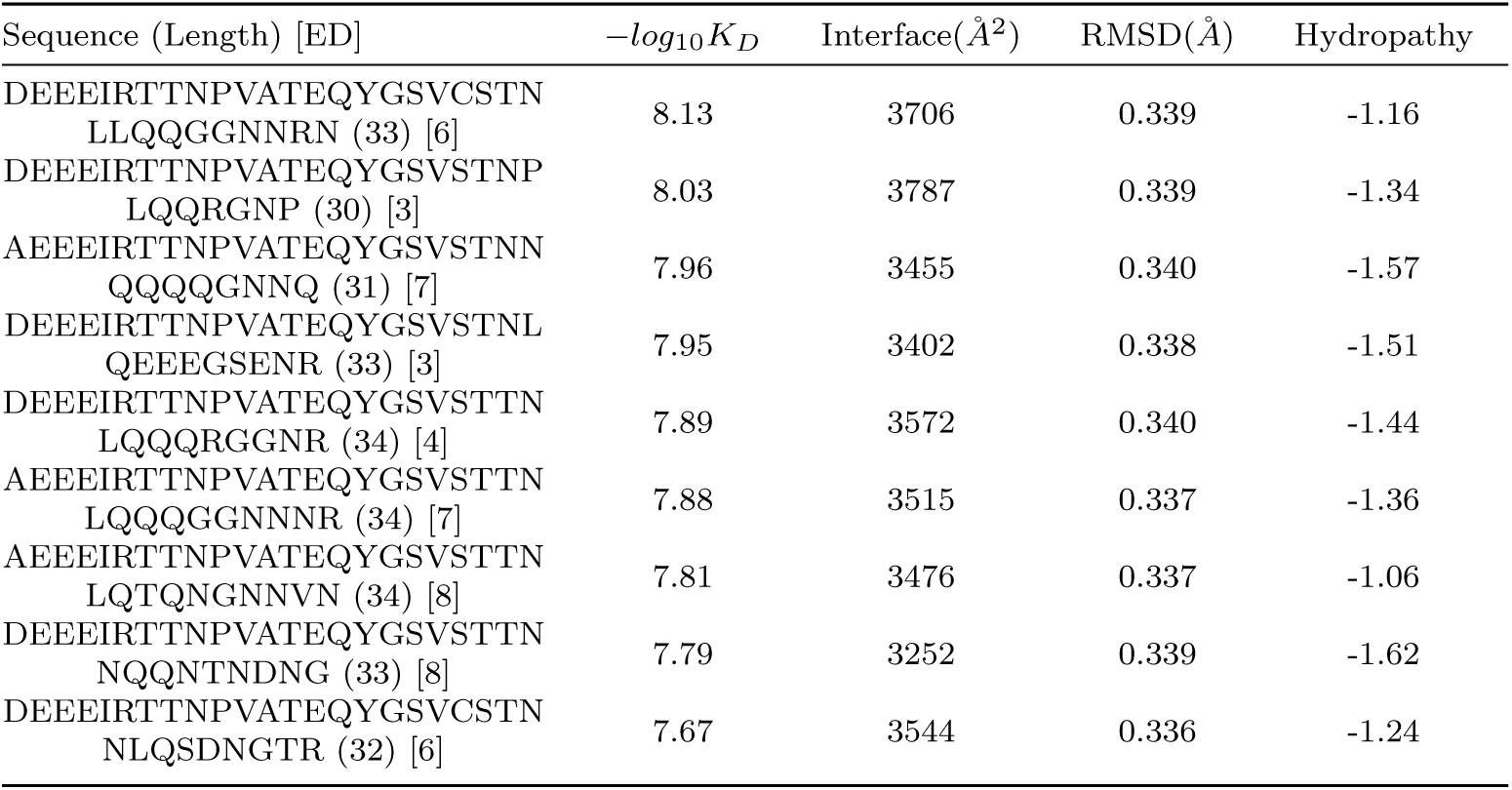
Properties of the top 9 candidates selected for hTfR1 Binding. This table presents the amino acid sequences of the top 9 hTfR1-binding candidates and their key properties, including sequence length, edit distance from the WT (ED), binding affinity to hTfR1 (*−log*_10_*K_D_*), interface area (Interface), structural deviation (RMSD), and grand average of hydropa-thy values (Hydropathy).

To investigate the ability of AAVDiffusion to capture the evolutionary sequence properties of Viable AAVs, we calculated Shannon entropy for each position in the MSA of Viable AAVs and AAVDs, with MSA constructed using Clustal Omega [53, 54]. Low Shannon entropy values represent highly conserved positions, while high entropy indicates high amino-acid diversity at a given position. The positional entropy in AAVDs is highly similar to that of Viable AAVs (Pearson correlation coefficient = 0.91*, P <* 1 10*^−^*^16^; *m.s.e.* = 0.033) (Fig. 4c). Even at position 25, the most variable site, the frequencies of individual amino acids in Viable AAVs and AAVDs are highly correlated (Pearson correlation coefficient = 0.80*, P <* 1 10*^−^*^4^; Sup. Fig. 1). Additionally, motif logos were constructed using Logomaker [55] to visualize amino acid preferences at each position. The logos indicate that AAVDs preserve the amino acid preferences observed in Viable AAVs, particularly at conserved positions (Fig. 4d). These results demonstrate that AAVDiffusion effectively reproduces the evolutionary sequence properties of Viable AAVs, capturing both sequence conservation and amino acid preferences.

To explore the capacity of AAVDiffusion to capture the specific local positional order of amino acids across Viable AAVs, we employed the *Z_m_* positional score function [56] to calculate amino-acid association for Viable AAVs and AAVDs. For each pair of the 20 amino acids, *Z_m_*(*a, b*) measures the average distance between amino acid *a* and the next occurrence of amino acid *b* in the sequence. The *Z_m_* scores for all amino acid pairs can be represented as a distance matrix (Fig. 4e). All *Z_m_* scores in the matrices for Viable AAVs and AAVDs are negative, suggesting that amino acids are closer than expected in random sequences with the same amino-acid frequency. On average, the positional order similarity between Viable AAVs and AAVDs is 91%, indicating that AAVDiffusion effectively captures the local amino-acid relationships of Viable AAVs.

To evaluate the similarity between the sequence spaces of AAVDs and Viable AAVs, we performed t-distributed stochastic neighbor embedding (t-SNE) [57] on their representations obtained from the embedding layer. The t-SNE visualization shows that AAVDs maintain a data manifold similar to Viable AAVs. However, differences in the distribution of dark dots, which indicate high data density, suggest that AAVDiffusion generates viable sequences without relying on memorization or repetition (Fig. 4f).

Finally, to assess the ability of AAVDiffusion to capture AAV structural features, we used SWISS-MODEL [58] to obtain the 3D structures of four sequences randomly selected from AAVDs. Then, we visualized and analyzed these structures using PyMOL [59]. The structures were aligned with the WT structure (PDB: 6IH9) [60], which was downloaded from the Protein Data Bank (PDB) [61]. The alignment reveals structural variations in the designed regions, with three sequences generated by AAVDiffusion displaying alpha-helical structures compared to the disordered coil structure of the WT (Fig. 4g and Sup. Fig. 2). Despite the structural diversity of AAVDiffusion-generated sequences, their structures are globally similar to WT, with RMSDs ranging from 0.071Å to 0.089Å. These results indicate that AAVDiffusion preserves critical structural features while enabling the generation of diverse and innovative sequences. Overall, these findings demonstrate that AAVDiffusion effectively learns the intrinsic relationships within viable AAV sequences, capturing critical features at both the sequence and structural levels. Moreover, it enhances diversity, enabling the design of novel AAV sequences.

### 2.5 Analysis of diversity in generated sequences across different positions

We further explore the correlates of sequence diversity at different positions in generated sequences with the 3D structure of the designed region, which encompasses buried, interface, and surface areas (Fig. 5a). To analyze the substitution and insertion diversity of AAVDs with viability probabilities *>* 0.5, we developed a dynamic programming algorithm to generate a substitution-insertion pattern for each sequence. This pattern defines the residue substitutions and insertions needed to transform the WT sequence into its corresponding generated sequence. Using the patterns generated for all sequences, we first counted the occurrences of each residue substitution and insertion at each position (Fig. 5b). Building on this, we calculated the mean number of substituted and inserted residue types at each position (Fig. 5c).

**Fig. 5.**
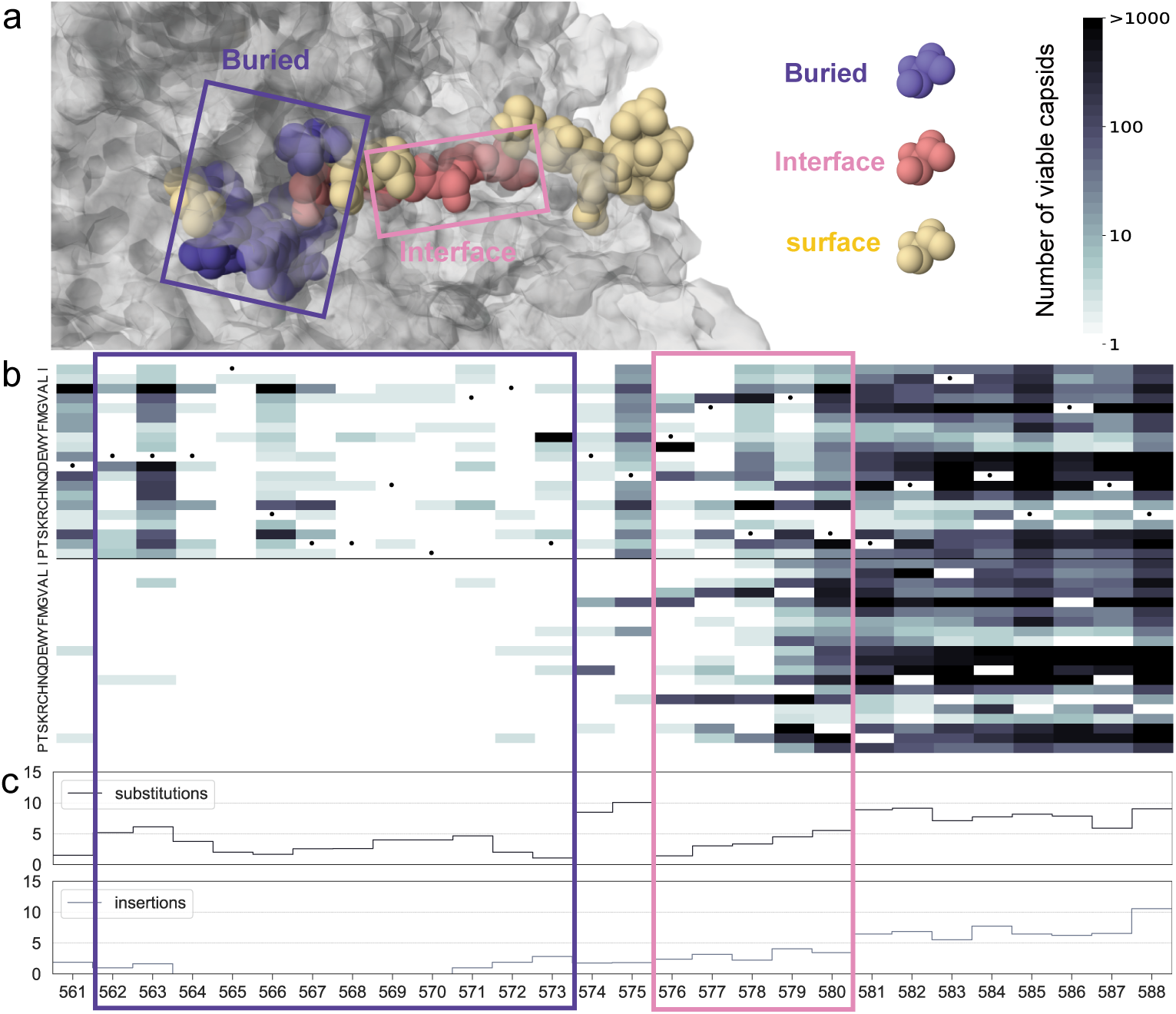
Sequence diversity in the selected AAVDs across the designed region and its structural context. **a**, 3D structure of the 28-residue designed region with buried (purple), inter-face (pink), and surface (yellow) regions, shown in context with interfacing monomers. **b**, Top: heatmaps showing substitutions within selected AAVDs. The dark dots mark the WT residues. Bot-tom: heatmaps showing insertions within selected AAVDs. **c**, Top: mean number of substituted residue types at each position across selected AAVDs. Bottom: mean number of inserted residue types at each position across selected AAVDs.

We observed that the first two-thirds of the designed region exhibit a strong preference for particular amino acids at each position (Fig. 5b), which is also reflected in the low mean number of residue types (Fig. 5c). This conservation is likely due to the reduced surface exposure of these positions (Fig. 5a) and the constraints imposed by the oligomeric interface [22]. Despite this conservation, AAVDiffusion successfully incorporated diverse residue substitutions at buried and interface sites, as well as introducing varied insertions at interface sites. Moreover, it also managed to incorporate insertions into the buried part of the capsid, an area typically intolerant to such modifications (Fig. 5b). These results demonstrate the capability of AAVDiffusion to understand the spatial constraints of the designed region while accommodating diverse sequence variations to meet high-viability requirements, without compromising structural integrity.

### 2.6 AAVDiffusion generates stable structures

Previous studies have shown that low-stability VP1 proteins are rapidly degraded by the proteasome, thereby preventing capsid assembly [62, 63]. Therefore, a critical aspect of designing AAV sequences is ensuring they fold into stable structures. To evaluate whether AAVDiffusion generates stable AAV sequences, we compared the stability of AAVDs with that of Viable AAVs. We first used SWISS-MODEL to generate the 3D structures of AAVDs and viable AAVs. We then performed Rosetta-RelaxBB runs [64] on these structures to calculate Rosetta Energy Units (REU) per residue, with lower values indicating greater protein stability [64, 65].

The mean REU per residue of AAVDs is –0.99, which is 0.02 lower than that of Vibale AAVs (Fig. 6a). This suggests that AAVDs are more stable than Viable AAVs. To identify residues contributing to enhanced stability, we selected the top 100 most stable AAVDs and generated a motif logo using WebLogo [66], based on the MSA of these sequences and the WT sequence. The logo reveals distinct amino acid preferences in the interface region, which differ from the WT and are likely to contribute to the increased stability (Fig. 6b). Based on this analysis, we selected the top 3 most stable AAVDs (numbered 5338, 9487, and 17768) and compared their 3D structures with the WT structure. The result shows that the interface region exhibits a coil structure in the WT, whereas the stable AAVDs display alpha-helical structures (Fig. 6c). The formation of these alpha-helical structures is likely a key factor contributing to the increased stability. All these results suggest that AAVDiffusion generates sequences with enhanced stability, possibly by introducing alpha-helical structures.

**Fig. 6.**
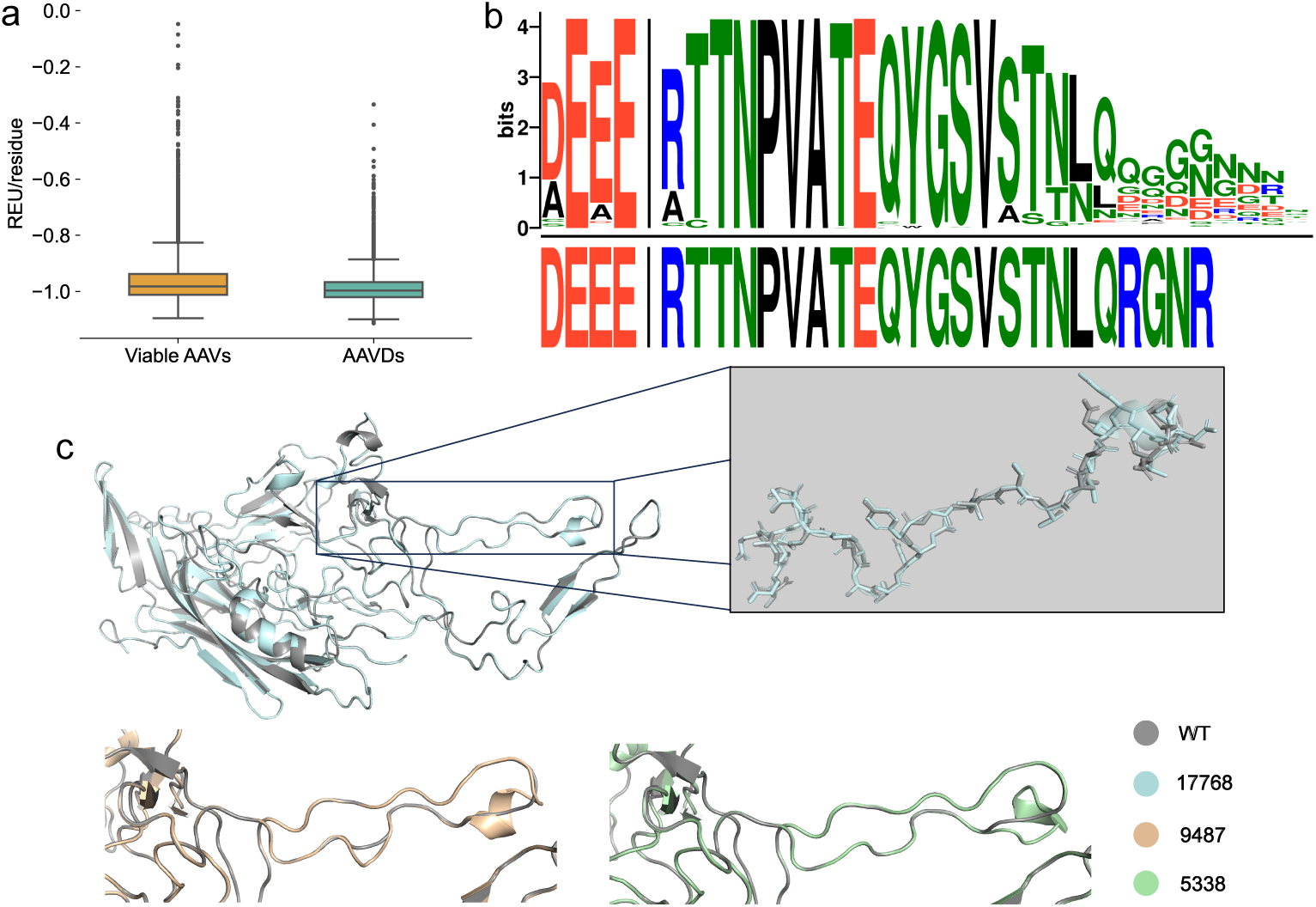
AAVDiffusion-generated sequences exhibit enhanced stability. **a**, Distributions of Rosetta Energy Units per residue for Viable AAVs and AAVDs. **b**, Sequence motif logo for the MSA of the 100 most stable AAVDs and the WT sequence, with the WT sequence shown at the bottom. **c**, Structural alignments of the top 3 most stable AAVDs (17768, 9487, and 5338) with the WT structure. The WT structure is shown in gray, and the AAVDs are shown in blue (17768), orange (9487), and green (5338). The designed region is magnified in the first alignment and displayed in stick mode to show structural details.

### 2.7 In silico Screening for hTfR1-Targeting AAV Capsids

Developing AAV vectors capable of effectively delivering genes throughout the human central nervous system (CNS) has the potential to significantly expand the range of treatable genetic diseases. AAVs can traverse the BBB by binding to hTfR1, a protein expressed on the BBB.

To identify promising hTfR1-targeting AAV candidates, we applied a selection workflow to AAVDs (Sup. Fig. 3). The workflow began by removing AAVDs already present in the AAV dataset. Next, we retained sequences with a viability probability predicted by AAVDiffison-classifier *>* 0.8. The retained sequences were ranked based on their REU per residue, and the top 3,000 most stable sequences were selected. We then evaluated their affinity for hTfR1 (denoted as − *log*_10_*K_D_* in Fig. 7a and Table 1) using HDOCK docking [67] and AREA-AFFINITY predictions [67], where higher values indicated stronger binding affinity. For affinity screening, we used BI-hTfR1 as a benchmark, an AAV capsid designed by Huang et al. that effectively crosses the BBB. Finally, sequences with affinity higher than that of BI-hTfR1 were retained, resulting in 196 candidates.

**Fig. 7.**
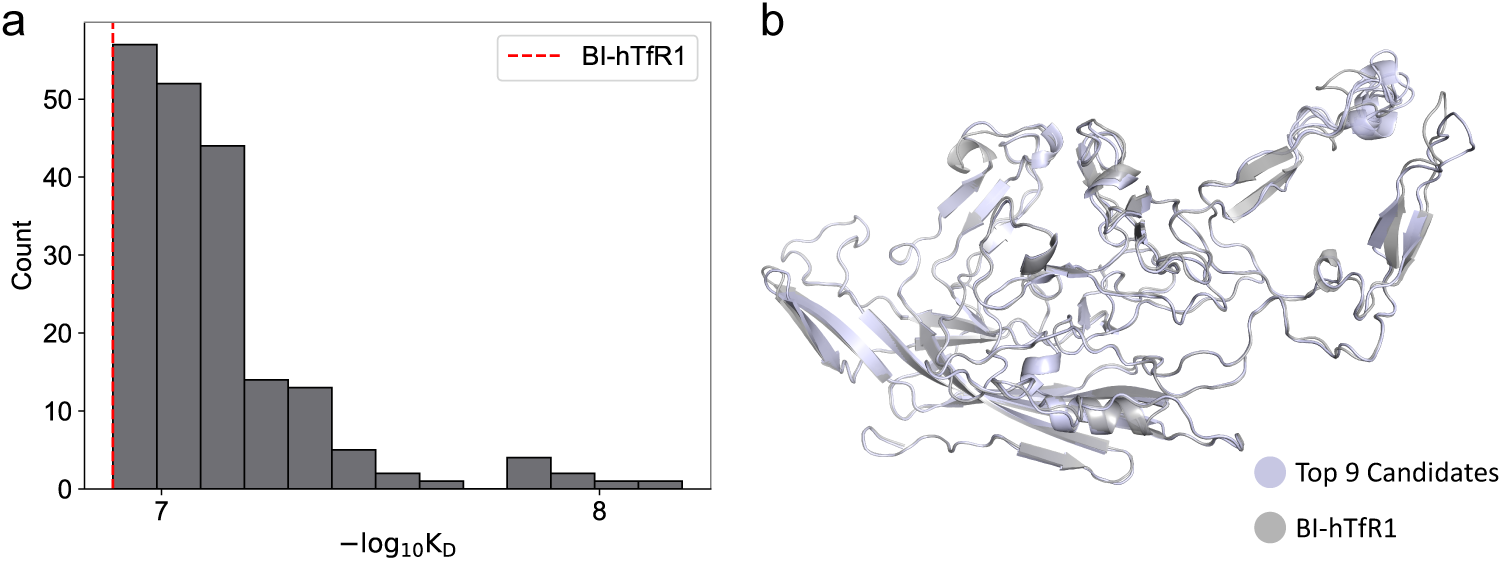
196 diverse hTfR1-targeting AAV capsids are identified through selection workflow. **a**, Binding affinity distribution of hTfR1-targeting candidates. The red dashed line represents the affinity of BI-hTfR1 (*−log*_10_*K_D_* = 6.89). **b**, Structural comparison of the top 9 candidates (light purple) with the BI-hTfR1 structure (gray).

The 196 candidates show a range of affinities for hTfR1, with 138 exhibiting affinities *>* 7.0 and 2 exceeding 8.0, significantly higher than the affinity of 6.89 for BI-hTfR1 (Fig. 7a). To further characterize these candidates, we analyzed the top 9 candidates with the highest affinities. These candidates vary in sequence length (ranging from 30 to 34 amino acids) and edit distance from the WT (ranging from 3 to 8 amino acids) (Table 1). Despite this sequence diversity, their structures are globally similar to BI-hTfR1, with all-atom VP1 RMSDs ranging from 0.336 Å to 0.340 Å (Fig. 7b and Table 1).

Although the overall structures are similar, each candidate exhibits a distinct binding mode with hTfR1 (Sup. Fig. 4 and Sup. Fig. 5). To explore the binding determinants, we calculated the surface area buried by each candidate when bound to hTfR1 (denoted as Interface in Table 1). Several candidates show larger binding interface areas than BI-hTfR1, which has an interface area of 3510 *Å*^2^. This suggests that these candidates may form novel interactions with hTfR1. Interestingly, no correlation was observed between binding interface area and affinity, indicating that surface area burial is not the primary determinant of binding. We also calculated the grand average of hydropathy values (GRAVY) [68] for each candidate to assess residue hydrophobicity (denoted as Hydropathy in Table 1). Similar to the interface area, hydropathy values vary but show no correlation with affinity, suggesting that hydrophobicity is not a major determinant of binding. Overall, these results indicate that the binding affinities of the candidates are primarily shaped by the sequence design of AAVDiffusion rather than dictated by a single, common binding mechanism.

## 3 Discussion

AAV-based gene delivery vectors are achieving increasing success in human clinical trials, offering the potential to treat a wide range of genetic and non-genetic disorders. However, the natural properties of AAV capsids significantly limit their efficacy. Although AAV capsid library selection and directed evolution have facilitated the development of improved vectors, existing methods for generating AAV capsid libraries often produce nonviable variants that fail to assemble or package DNA properly. In this study, we developed AAVDiffusion, a viability-guided diffusion model designed for de novo generation of viable AAV capsid protein sequences. AAVDiffusion employs a reverse diffusion process to transform noise distribution into the desired AAV sequence distribution, thereby generating viable AAV sequences. To better capture longrange dependencies and structural information within AAV sequences, we utilized a Transformer-based architecture for denoising. Additionally, to enhance its understanding of functional peptides and improve sequence diversity, we pre-trained the diffusion module on peptide sequences from the UniProt database. To further optimize viability, we incorporated a viability classifier that evaluates sequence viability based on latent representations and enables gradient-based viability optimization during generation.

Extensive evaluation of AAV candidates demonstrates that AAVDiffusion significantly outperforms existing methods in generating viable AAV sequences, achieving the highest median viability probability across all classifiers and surpassing the second-best model by 0.19 under CNN(C1+R2) evaluation. Through a series of comprehensive analyses, including Pearson correlation between predicted viability probabilities and experimental production scores, comparison of amino acid composition, assessment of Shannon entropy, evaluation of *Z_m_* scores, t-SNE visualization of sequence space, and 3D structural analysis, we demonstrate that AAVDiffusion effectively captures the intrinsic relationships within viable AAV sequences. Furthermore, our selection workflow identified 196 AAV candidates with higher binding affinities to hTfR1 than BI-hTfR1, suggesting their potential to cross the BBB more efficiently and serve as promising vectors for targeted CNS delivery.

## 4 Methods

### 4.1 Data preparation

Our training dataset consists of two main components: the AAV dataset for fine-tuning and the UniProt dataset for pre-training. The AAV dataset is constructed by merging three datasets from Bryant et al. [22], comprising 296,913 sequences. Among these, 153,692 sequences are viable and 143,221 sequences are nonviable, with sequence lengths ranging from 28 to 43 amino acids. Each sequence is accompanied by an experimental production score and a binary viability label. The experimental production score, derived from high-throughput experiments, evaluates the production efficiency of AAV sequences. A higher score indicates a greater likelihood of successful capsid assembly and genome packaging. The binary viability label is determined based on the production score. It reflects the production viability of AAV sequences, where “1” represents viability and “0” represents nonviability.

The UniProt dataset is sourced from the UniProt-SwissProt and UniProt-Trembl databases [44]. For consistency with the AAV dataset, we retained sequences ranging from 28 to 43 amino acids in length. Sequences containing non-natural amino acids (B, J, O, U, X, Z) or lowercase letters were then excluded. After filtering, over 15 million sequences remained, of which 3 million were randomly sampled for pre-training.

Furthermore, to train and evaluate the AAV sequence diffusion and viability classifier modules, both the UniProt and AAV datasets were divided into training, validation, and test sets in an 8:1:1 ratio.

### 4.2 AAVDiffusion

Recently, continuous diffusion models have been adapted for discrete text data, achieving remarkable performance [69, 70]. Inspired by the analogies between protein sequences and human languages [71, 72], we propose AAVDiffusion, a model that integrates a continuous diffusion model with a viability classifier for the generation of viable AAV capsid sequences. AAVDiffusion consists of two major modules, including AAV sequence diffusion module and the viability classifier module. In the following sections, we detail these modules and explain how their integration enables the generation of high-viability AAV sequences.

#### AAV sequence diffusion module

The AAV sequence diffusion module employs a continuous diffusion model to generate diverse AAV sequences. We define an AAV capsid sequence of length *n* as *s* = (*s*_1_*, …, s_n_*). Since standard continuous diffusion models cannot be directly applied to discrete AAV sequence data, we introduce a trainable embedding function *E*(*s_i_*) to map each amino acid *s_i_* to a latent space. The embedded representation of the sequence *s* is then *E*(*s*) = [*E*(*s*_1_)*, …, E*(*s_n_*)]. To adapt discrete sequence input into the standard forward process, we define a Markov transition *q_ϕ_* (*z*_0_|*s*) = *N* (*E* (*s*) *, β*_0_*I*) to map the sequence *s* to its latent representation *z*_0_. This representation not only captures key sequence features but also generates a more compact and smooth latent space, enhancing the training efficiency of the diffusion model [73]. The embedding function is implemented as a standard neural network embedding layer with a weight matrix of size 20 by 16, where 20 corresponds to the number of naturally occurring amino acids in the input space and 16 represents the latent space dimensionality.

To decode *z*_0_ back to a discrete sequence *s*, we introduce a trainable decoding function parameterized as 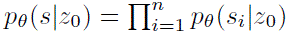, where *p_θ_*(*s_i_*|*z*_0_) represents a softmax distribution over all possible amino acids at position *i*. This function is implemented using a fully connected layer, which computes the probability distribution of amino acids at every sequence position. The final sequence is generated by selecting the most probable amino acid at each position via *argmax p_θ_* (*s z*_0_).

With our embedding and decoding functions, we can represent AAV sequences using lower-dimensional latent representation. This latent representation is then processed by a latent diffusion model, which consists of a diffusion process and a denoising process to generate a denoising latent representation *z*_0_. In the diffusion process, we treat the latent representation of the AAV sequence as a set of particles evolving in a thermodynamic system [74, 75]. Over time, the latent representation experiences gradual corruption with Gaussian noise *ɛ*, ultimately transitioning into pure Gaussian noise at diffusion step *T*. The diffusion process follows a Markov chain [76]:

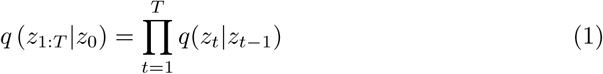

where *q*(*z_t_ z_t−_*_1_) represents the Markov diffusion kernel, which applies noise to the latent representation at each time step. Following the formulation in [42], we define the diffusion kernel as:

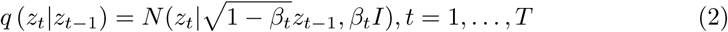

where *β*_1_*, …, β_T_* are variance schedule hyperparameters controlling the noise level at each diffusion step.

The generative process is modeled as the reverse of the diffusion process, starting from the latent representation *z_T_* sampled from the distribution *N* (0*, I*). This latent representation is iteratively refined through a learned denoising process to reconstruct the desired latent representation *z*_0_. We model the denoising process as a Markov chain with learnable transitions:

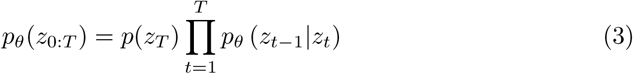

To ensure that the denoised *z*_0_ is precisely centered at an amino acid embedding, we employ a denoising model *f_θ_* (*z_t_, t*) to explicitly estimate *z*_0_. This estimate serves as the objective during training. At decoding time, we also apply the clamping trick from DiffusionLM [70]. Spesifically, AAVDiffusion denoises *z_t_* to *z_t−_*_1_ by first computing an estimate of *z*_0_ via *f_θ_* (*z_t_, t*). The predicted *z*_0_ is then mapped to its nearest protein sequence embedding through the decoding and encoding layers. Finally, *z_t−_*_1_ is sampled conditioned on this embedding. Thus, the transformation from *z_t_* to *z_t−_*_1_ is formulated as follows:

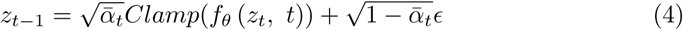

where 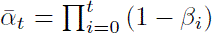 and *ɛ*∼*N* (0, 1). The clamping trick forces the predicted *z*_0_ to commit to an amino acid for intermediate diffusion steps, thereby improving prediction precision and reducing rounding errors.

The denoising model *f_θ_* is implemented using a Transformer-based architecture with 12 layers of Transformer encoders, utilizing the self-attention mechanism as described by Vaswani et al. [27]. Temporal information is incorporated using time step embedding, similar to positional embedding.

The maximum sequence length of the AAV sequence diffusion module is 43, with 2000 diffusion steps and a square-root noise schedule. End-to-end training is performed on the embedding layer, denoising model, and decoding layer. The loss function is defined as:

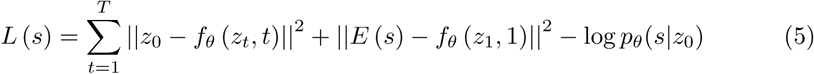

where *L*(*s*) consists of three terms. The first term, 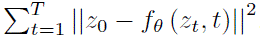, measures the error between the predicted latent representation *f_θ_* (*z_t_, t*) and the original latent representation *z*_0_, promoting accurate denoising at each step. The second term, 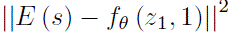, enforces consistency between the output of the embedding layer and the predicted latent representation *f_θ_* (*z*_1_, 1), encouraging the embedding layer to capture meaningful structural and functional information of AAV sequence. The final term, −log *p_θ_*(*s*|*z*_0_), maximizes the likelihood of the target AAV sequence *s* given the denoised latent state *z*_0_, encouraging the generation of realistic and accurate AAV sequences.

To enhance the ability of the diffusion module to capture the complex characteristics of functional peptides and generate more diverse sequences, we do not restrict the AAV sequence diffusion module to learning only the density of the AAV dataset. Instead, we pre-train the module on the UniProt dataset and then finetune it on the AAV dataset. Pre-training is conducted using the Adam optimizer with a learning rate of 0.0001, a dropout rate of 0.1, a batch size of 64, and 200,000 training iterations. Fine-tuning is performed for 15,000 iterations with all other hyperparameters remaining unchanged.

#### Viability classifier module

To improve the viability of AAVDiffusion-generated sequences, we introduce the viability classifier module, which evaluates sequence viability based on latent representations and enables gradient-based viability optimization during sequence generation.

The viability classifier module is constructed through the following steps. First, each AAV sequence *s* with a viability label *a* is transformed into its latent representation *z*_0_ via *q_ϕ_* (*z*_0_|*s*) = *N* (*E* (*s*) *, β*_0_*I*). Subsequently, a diffusion process is applied to generate a series of latent representations *z*_1:_*_T_*. Together with *z*_0_, these representations form the complete set of latent vectors *z*_0:_*_T_*. Next, the time step *t* is embedded to match the dimension of *z_t_*, and this embedding is concatenated with both the positional embedding and *z_t_* to form the final input to the classifier. Finally, the classifier *p_ξ_*(*a z_t_*) is trained on these inputs to predict the viability.

To implement this classifier, we adopt a fully connected neural network architecture consisting of two fully connected layers, with a Rectified Linear Unit (ReLU) activation [77] between them, and a final softmax layer for categorical prediction. It is trained using the binary cross-entropy loss to minimize the discrepancy between predicted and true viability labels. The Adam optimizer is employed for parameter optimization with a learning rate of 0.0001, a batch size of 256, and 300 training epochs.

For viability control, we employ gradient backpropagation from the classifier outputs to the latent inputs *z*_0:_*_T_*. This process is equivalent to decoding from the posterior 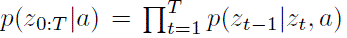, which can be decomposed into a sequence of control problems at each diffusion step: *p*(*z_t−_*_1_ *z_t_, a*) ∝ *p*(*z_t−_*_1_ *z_t_*) *p*(*a z_t−_*_1_*, z_t_*). Following conditional independence assumptions from prior work on controlling diffusions [78], we simplify *p*(*a*|*z_t−_*_1_*, z_t_*) = *p*(*a*|*z_t−_*_1_). Thus, the gradient update for *z_t−_*_1_ is expressed as:

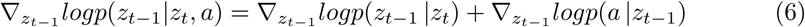

Here, *logp*(*z_t−_*_1_ |*z_t_*) is parameterized by the AAV sequence diffusion module, while *logp*(*a z_t−_*_1_) is parameterized by the viability classifier module. Both terms are differentiable.

Unlike controllable generation methods for diffusions that do not include *logp*(*z_t−_*_1_ |*z_t_*) in the objective [79, 80], AAVDiffusion performs gradient updates on a control objective with regularization: *λlogp*(*z_t−_*_1_ *z_t_*) + *logp*(*a z_t−_*_1_), where *λ* is a hyperparameter balancing high AAV sequence likelihood and high viability. This regularization helps prevent excessive control, which could lead to unrealistic AAV sequences. A grid search identifies *λ* = 0.0005 as the optimal value.

To further enhance control quality, we take multiple gradient steps for each diffusion step. The control intensity is governed by two parameters: *k* (the number of gradient steps per diffusion step) and *η* (the learning rate per gradient step). The grid search also reveals the optimal configuration: three Adagrad updates per diffusion step (*k* = 3) with a learning rate of 0.01 (*η* = 0.01).

To mitigate the computational cost introduced by multiple gradient steps, we employed DDIM [81], reducing the number of diffusion steps from 2000 to 50, significantly accelerating sequence generation while preserving sample quality.

### 4.3 Compared methods

We compared the performance of AAVDiffusion in generating viable AAV sequences with four methods. These methods include ClaSS [37], MSA-VAE [34], ProteinGAN [39] and DeepAAV [22].

#### ClaSS

ClaSS is a method that combines a deep generative autoencoder with classifiers to achieve controlled protein sequence generation [37]. It first trains a global deep autoencoder over all known short peptide sequences derived from various organisms, including target protein sequences and sequences from the UniProt database. The target protein sequences are then mapped into the autoencoder’s latent space to obtain their latent representations. A density model and multiple attribute classifiers are subsequently constructed based on these representations. During the controlled generation of target proteins, latent vectors are sampled from the density model and evaluated by the classifiers for their likelihood of exhibiting the desired properties. Then, a rejection sampling strategy is applied to filter the latent vectors based on their predicted properties, and the selected ones are decoded into protein sequences. To reproduce ClaSS, we trained the model from the GitHub repository using our UniProt dataset and AAV dataset, following the tutorial provided by the authors: https://github.com/IBM/controlled-peptide-generation.

#### MSA-VAE

MSA-VAE is a variational autoencoder designed to model the MSA of a target protein family, enabling the generation of novel sequence variants [34]. It takes as input the rows of the MSA, retaining only the columns corresponding to positions in the target protein. The model employs a fully connected feed-forward encoder-decoder architecture to learn the patterns of sequence variation that underlie function. To reproduce MSA-VAE, we first aligned the AAV dataset using Clustal Omega [53, 54] and retained only the MSA columns corresponding to positions in the WT sequence. The model was then trained on this processed MSA input, and novel AAV sequence variants were generated following the tutorial provided by the authors: https://github.com/alex-hh/deepprotein-generation.

#### ProteinGAN

ProteinGAN is a self-attention-based variant of GANs designed for generating novel functional protein sequences with natural-like biochemical properties [39]. Its architecture consists of a discriminator and a generator, both incorporating ResNet blocks and a self-attention layer to capture long-range dependencies in protein sequences. Through adversarial training on functional protein sequences, the generator learns the underlying amino acid relationships, enabling it to transform sampled noise into “real” functional protein sequences. To reproduce ProteinGAN, we first applied its preprocessing method to assign dynamic up-sampling weights to the AAV dataset, ensuring a balanced distribution. We then trained the model on this processed dataset and generated AAV sequence variants by following the tutorial provided by the authors: https://github.com/Biomatter-Designs/ProteinGANprovidedbytheauthors.

#### DeepAAV

DeepAAV is a method for generating viable AAV capsid proteins using viability classifiers [22]. It constructs nine classifiers by training three different model architectures (CNN, LR, and RNN) on three distinct datasets (C1+R2, C1+R10, and R10+A39) to estimate the viability of AAV sequences. For each classifier, the method iteratively mutates the WT sequence, ranks and filters the variants over 20 rounds to generate the final model-designed sequences. For comparison with AAVD-iffusion, we collected six datasets generated by three CNN classifiers and three RNN classifiers, including CNN(C1+R2), CNN(C1+R10), CNN(R10+A39), RNN(C1+R2), RNN(C1+R10) and RNN(R10+A39). These datasets are available in the GitHub repository https://github.com/alibashir/aav. From each dataset, we sampled 19,680 sequences and evaluated their viability using the classifier that designed them.

### 4.4 Evaluation metric and bioinformatic analyses

In this section, we introduce the evaluation metric and bioinformatic analyses employed in this study.

#### Viability prediction

The viability of AAV sequence variants was assessed using the three CNN and three RNN classifiers mentioned above. Each classifier predicts a viability probability (*P* ∈ [0, 1]), with higher values indicating a greater probability of a sequence being viable for packaging of a DNA payload. These classifiers were run following the tutorial provided by the authors: https://github.com/google-research/google-research/tree/master/aav.

#### Multiple sequence alignment

All MSAs in this study were constructed using Clustal Omega v1.2.4 [53, 54] on a Linux-based system.

#### Shannon entropy calculation

Shannon entropies were calculated based on the MSA of Viable AAVs and AAVDs. First, Viable AAVs and AAVDs were combined and aligned to construct a global MSA. This alignment was then separated into two datasets corresponding to Viable AAVs and AAVDs. Columns with more than 90% of gaps in either dataset were excluded from further analysis. Finally, Shannon entropy was calculated for each remaining column in the MSA:

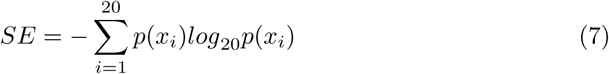

where *p*(*x_i_*) represents the frequency of amino acid *i* at a given column in the MSA.

#### *Z_m_* positional score calculation

The *Z_m_* positional score [56] was calculated for every possible amino acid pair (*a, b*) in a sequence and averaged over the whole dataset. The *Z_m_* positional score function is defined as:

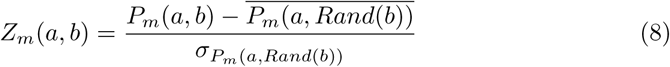

where 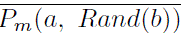 and 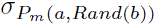 represent the mean and standard deviation of the randomly shuffled sequence association for the amino acid pair.

The association function used for scoring was the minimal proximity function:

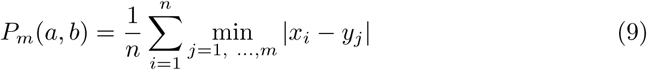

where for each position *x_i_* of amino acid *a*, the closest occurrence of amino acid *b* at position *y_j_* is identified, and the average distance between the pairs is computed.

#### t-SNE plot generation

The embedded representations of Viable AAVs and AAVDs from the embedding layer were used as input for t-SNE. The t-SNE module from scikit-learn v1.1.1 [82], a Python library, was applied with default settings (perplexity 30, early exaggeration 12, learning rate 200, maximum of 1,000 iterations). The resulting t-SNE coordinates were visualized using seaborn v0.12.2 [83], a Python data visualization library.

#### Protein 3D structure visualization and analysis

All protein 3D structure visualizations, as well as RMSD and binding interface area calculations, were performed using PyMOL [59].

#### Protein 3D structure prediction

All predicted protein structures in this study were generated using SWISS-MODEL [58], a homology modeling server avail-able at https://swissmodel.expasy.org. To predict the 3D structures of complete VP1 sequences corresponding to Viable AAVs and AAVDs, each Viable AAV and AAVD sequence was embedded into the VP1 template. The resulting complete VP1 sequences, along with the WT VP1 protein structure (PDB: 6IH9) [60] downloaded from the PDB, were uploaded to SWISS-MODEL for structure prediction using the user-template mode.

#### Substitution-insertion pattern generation

To analyze the diversity of insertions and mutations at each position within the AAVDs, we developed a dynamic programming algorithm to generate one possible substitution-insertion pattern for each AAVD. This pattern follows the substitution-insertion pattern defined by Bryant et al. [22], which consists of 28 substitution positions (i.e., WT position) and 28 insertion positions. For each substitution position, any of the 19 amino acids different from WT are allowed, while for each insertion position, all 20 standard amino acids are considered. Residue substitutions and insertions, as defined in this pattern, enable the transformation of the WT sequence into its corresponding sequence variant.

#### REU per residue calculation

REU per residue was calculated using the Rosetta Relax protocol, implemented in the Rosetta Software Suite v2024.09 [64, 65]. The input structures were VP1 protein structures corresponding to Viable AAVs and AAVDs, predicted by SWISS-MODEL. Each structure underwent a single replica run with five relaxation iterations.

#### Motif logos construction

Motif logos in this study were generated using two tools. For the MSA of Viable AAVs and AAVDs, motif logos were constructed using Logomaker v0.8 [55] after excluding columns with more than 90% gaps. Additionally, WebLogo [66] (https://weblogo.berkeley.edu) was used to analyze the top 100 AAVDs ranked by stability, which were aligned to the WT sequence while maintaining all alignment positions.

#### Docking of VP1 proteins with hTfR1

The Linux version of HDOCK [84] was used to generate docking complexes between the VP1 protein structures of AAVDs and hTfR1. The hTfR1 structure (PDB: 1SUV) was obtained from PDB. Similarly, HDOCK was employed to predict the docking complex between BI-hTfR1 and hTfR1. The BI-hTfR1 structure was predicted using SWISS-MODEL based on its amino acid sequence.

#### Binding affinity prediction

All binding affinity predictions in this study were performed using AREA-AFFINITY [67] (https://affinity.cuhk.edu.cn), a web server for protein-protein binding affinity prediction, with docking complexes generated by HDOCK as inputs.

